# Seasonal recurrence and modular assembly of an Arctic pelagic marine microbiome

**DOI:** 10.1101/2024.05.10.593482

**Authors:** Taylor Priest, Ellen Oldenburg, Ovidiu Popa, Bledina Dede, Katja Metfies, Wilken-Jon von Appen, Sinhué Torres-Valdés, Christina Bienhold, Bernhard M. Fuchs, Rudolf Amann, Antje Boetius, Matthias Wietz

## Abstract

Deciphering how microbial communities are shaped by environmental variability is fundamental for understanding the structure and function of ocean ecosystems. Thus far, we know little about the structuring of community functionality and the coupling between taxonomy and function over seasonal environmental gradients. To address this, we employed autonomous sampling devices and *in situ* sensors to investigate the taxonomic and functional dynamics of a pelagic Arctic Ocean microbiome over a four-year period. We demonstrate that the dominant prokaryotic and microeukaryotic populations exhibit recurrent, unimodal fluctuations each year, with community gene content following the same trend. The recurrent dynamics within the prokaryotic microbiome are structured into five temporal modules that represent distinct ecological states, characterised by unique taxonomic and metabolic signatures and connections to specific microeukaryotic populations and oceanographic conditions. For instance, *Cand*. Nitrosopumilus and the machinery to oxidise ammonia and reduce nitrite are signatures of early polar night, along with Radiolarians. In contrast, late summer is characterised by *Amylibacter*, sulfur compound metabolism and diverse Haptophyta lineages. Exploring the composition of modules further along with their degree of functional redundancy and the structuring of genetic diversity within functions over time revealed seasonal heterogeneity in environmental selection processes. In particular, we observe strong selection pressure on a functional level in spring while late polar night features weaker selection pressure that likely acts on an organismal level. By integrating taxonomic, functional, and environmental information, our study provides fundamental insights into how microbiomes are structured under pronounced environmental variability in understudied, yet rapidly changing polar marine ecosystems.

## INTRODUCTION

Bacteria, archaea and microeukaryotes are the dominant life forms in ocean environments and comprise an immense taxonomic, functional and physiological diversity. These microbes drive and respond to changes in their surrounding environment, such as bottom-up (e.g. resource availability) and top-down (e.g. grazing and viral infection) factors and physicochemical conditions (e.g. temperature), which results in the assembly of distinct communities over spatial and temporal scales^1,2^. The assembled communities subsequently perform essential trophic roles and mediate the biogeochemical cycling of biologically important elements^3–5^. Deciphering how microbial community dynamics are shaped across environmental gradients is thus fundamental for understanding the structure and function of ecosystems and how they respond to change.

Long-term observations have uncovered recurrent and transient dynamics in microbial communities across daily, seasonal and annual timescales. In temperate ecosystems, communities are predominantly structured by seasonal variability, with broadly recurrent fluctuations of taxa on annual scales^6–11^. These patterns have led to the conclusion that microbial responses to biological and environmental shifts are predictable^12^. However, recent evidence indicates that microeukaryotes exhibit weaker temporal structuring than prokaryotes^13^, suggesting different controlling mechanisms. Furthermore, high-frequency sampling has shown that population dynamics are highly ephemeral in nature, undergoing rapid, short-lived fluctuations that transpire over days^14–16^. Thus, while microbial communities are structured over time, they also undergo constant flux, reflecting the dynamic nature of ocean ecosystems.

Despite the wealth of knowledge gained from long-term observations, the focus primarily on taxonomic dynamics and on temperate and sub-tropical ecosystems has left many questions unanswered. In particular, it remains unclear how compositional shifts across environmental gradients translate to changes in the functionality of microbial communities. Since distantly related organisms can perform similar metabolic functions^17,18^, taxonomic information alone does not inform about ecological landscapes or ecosystem function. Therefore, long-term observations that integrate taxonomic, functional, and environmental information are greatly needed. This is particularly important in the polar oceans, where long-term observations are rare and unprecedented changes are taking place because of climate warming.

To address this, we investigated the dynamics of prokaryotic (here used operationally to refer to bacteria and archaea) and microeukaryotic communities from a taxonomic and functional perspective over a four-year period in an Arctic Ocean ecosystem, the West Spitsbergen Current (WSC). The WSC constitutes the primary entry route for Atlantic water into the Arctic Ocean and is characterised by pronounced seasonal variability in environmental conditions, archetypal of high-latitude ecosystems. However, unlike other Arctic Ocean ecosystems, the WSC remains ice-free year-round, due to the warmth of the North Atlantic water. Therefore, the WSC represents a model system for investigating the dynamics of microorganisms and ecosystem function in the context of pronounced seasonal variability. In addition, it can provide valuable insights into potential future shifts in Arctic Ocean ecosystems, given the progressive northward expansion of Atlantic water influence^19^.

We hypothesised that the Arctic epipelagic microbiome shows a seasonally recurring assembly primarily structured by polar day/night cycles. To investigate this, we employed moorings fitted with autonomous sampling and measuring devices to continuously track taxonomic, functional and environmental dynamics. Through 16S and 18S rRNA gene amplicon and PacBio HiFi metagenome sequencing, we generated a high-quality, temporally resolved microbiome catalogue. Using a Fourier transformation-based approach, we demonstrate that prokaryotic and microeukaryotic communities exhibit annually recurrent, seasonally structured dynamics. Within the prokaryotic microbiome, these dynamics assemble into five distinct temporal modules that feature unique taxonomic and metabolic signatures, are associated with specific microeukaryotic populations and are subject to different environmental selection pressures. Our study provides the first multi-year ecosystem catalogue from the Arctic that integrates taxonomic, functional and environmental information, and provides fundamental insights into the dynamics and structuring of microbiomes across pronounced environmental gradients.

## RESULTS & DISCUSSION

### The West Spitsbergen Current harbours an ecosystem with pronounced temporal structuring

We first investigated how environmental conditions in the WSC are structured over intra- and inter-annual scales. For this, we combined data collected from *in situ* sensors attached to the mooring, including temperature, salinity and oxygen saturation, with chlorophyll *a* measurements and satellite-derived values of photosynthetically active radiation (PAR) (Supplementary Table S1). Our measurements were derived from the epipelagic layer but varied between 20 – 100 m in depth due to the movement of the mooring by currents. Owing to the inclusion of multiple CTD sensors at different depths, we were also able to determine the lower bound of the mixed layer depth (MLD). Each annual cycle was characterised by pronounced shifts in environmental conditions (Figure 1). As expected, these shifts followed the transition between polar night and polar day, which is a major force stimulating biological dynamics in the Arctic^20^. The end of polar night was marked by an increase in PAR in April, which continued to rise until a maximum, on average, of 38 μmol photons m^−2^ s^−1^ in June. The increasing solar radiation onset warming, with temperatures rising until a peak of ∼7°C in August/September before decreasing again to <4 °C between December – May. Changes in MLD were inversely related to temperature (Pearson correlation: *R* = −0.47, *p* < 0.05), but were characterised by abrupt events of shallowing to <5 m in June and deepening to >200 m between December-January. Chlorophyll concentrations showed a lagged association with PAR, peaking between June-August. However, the magnitude and timing of the chlorophyll peak varied across years, from 3.35 μg l^−1^ in July 2019 to 13.22 μg l^−1^ in June 2018, indicating differences in phytoplankton bloom phenologies. The WSC thus exhibits pronounced intra-annual shifts in environmental conditions, presenting an ideal ecosystem to study seasonally driven biological dynamics.

**Figure 1.**
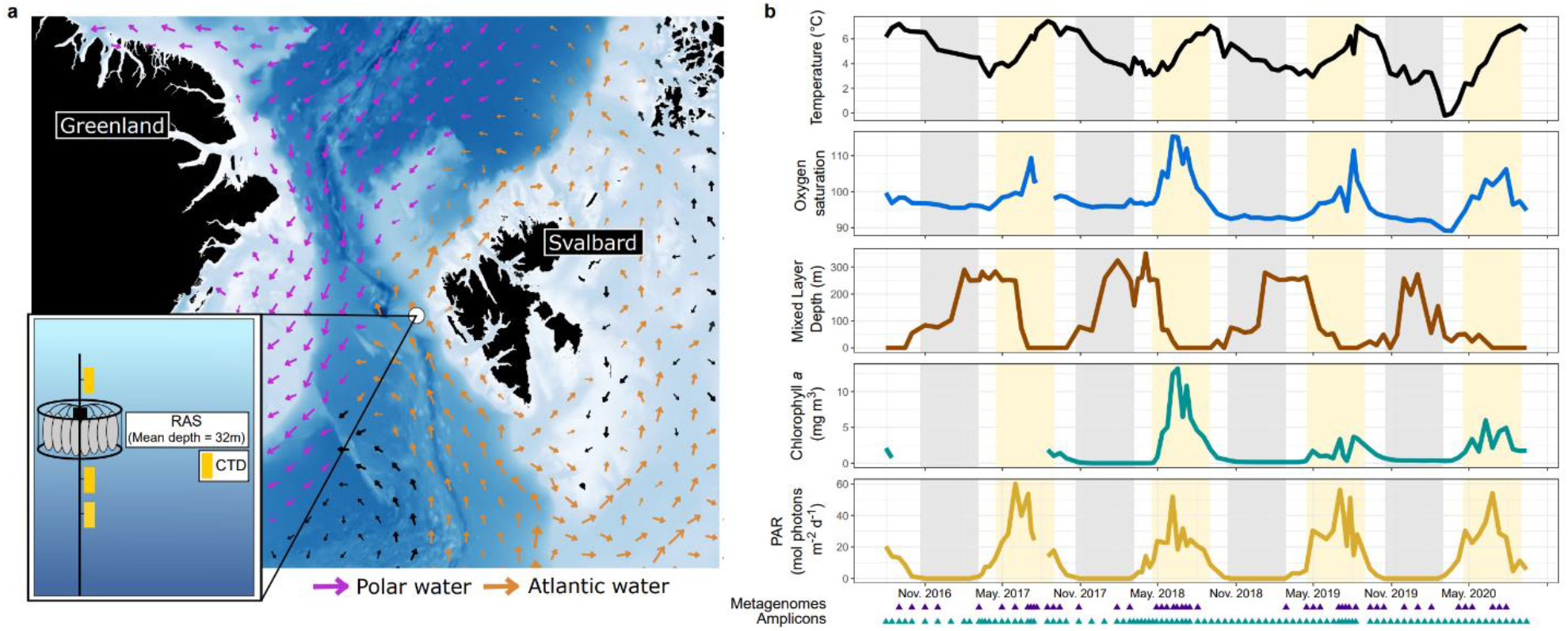
The WSC mooring site and environmental conditions between July 2016 and July 2020. **a)** Bathymetric map of the Fram Strait region with the mooring location within the West Spitsbergen Current. Arrows represent average current velocities over the four-year sampling period at the average depth of the moored autonomous sampler (32 m). **b)** Water temperature, oxygen saturation, mixed layer depth, chlorophyll concentrations measured from mooring-attached sensors and photosynthetically active radiation (PAR) derived from AQUA-MODIS satellite data. The shaded grey and yellow represent the periods of polar night and day, respectively.

### The intra- and inter-annual temporal structuring of communities

We next investigated the temporal structuring of prokaryotic and microeukaryotic communities from a taxonomic perspective. Using autonomous Remote Access Samplers, we collected 97 samples at, on average, fortnightly resolution that were used for 16S and 18S rRNA gene sequencing. From these, we recovered 3629 bacterial, 119 archaeal and 3019 microeukaryotic Amplicon Sequence Variants (ASVs) (Supplementary Table S2 and S3).

The alpha diversity of prokaryotic and microeukaryotic communities exhibited distinct trends within each annual cycle (Figure 2a and 2b). For prokaryotic communities, we observed a tight coupling between Species Richness (*R*), Evenness (*E*) and Shannon Diversity (*H’*), evidenced through significant positive Pearson’s correlations (Figure 2c and Supplementary Table S4 and S5). The alpha diversity metrics followed a unimodal fluctuation within each annual cycle for prokaryotic communities, reaching a peak during polar night (December - March) with a mean *R* of 1210 ± 208, *E* of 0.78 ± 0.04 and *H’* of 5.3 ± 0.36. The enriched diversity in winter observed here mirrors previous observations in temperate and polar regions^21–23^ and is associated with the deepening of MLD, which drives mixing and dilution of previously stratified communities^24^. The influx of PAR and rapid shallowing of MLD at the end of polar night coincided with a sharp drop in alpha diversity, with lowest values observed in June (*R* = 762 ± 156, *H’* = 4.50 ± 0.34, *E* = 0.68 ± 0.03). Hence, shifts in prokaryotic alpha diversity were tightly coupled to MLD, supported by strong positive Pearson’s correlations; Richness (*r* = 0.71, 95% CI [0.59,0.79], *p* = 4 × 10^−16^), Evenness (*r* = 0.43, 95% CI [0.25,0.58], *p* = 1.08 × 10^−5^) and Shannon diversity (*r* = 0.58, 95% CI [0.43,0.70], *p* = 5.68 × 10^−10^) (Supplementary Table S5). This observation is in contrast to previous reports of temperature^25–27^, ocean currents^28^ and day length^10,22^ as key drivers of epipelagic bacterial diversity. The greater role of MLD in shaping prokaryotic diversity observed here may be a feature unique to high-latitude ocean ecosystems, where seasonal shifts in MLD are more pronounced^29^ and can be influenced by sea-ice dynamics^30^. However, MLD has rarely been measured or incorporated before, and thus its considerations in future studies will help to ascertain whether its influence varies across latitude.

**Figure 2.**
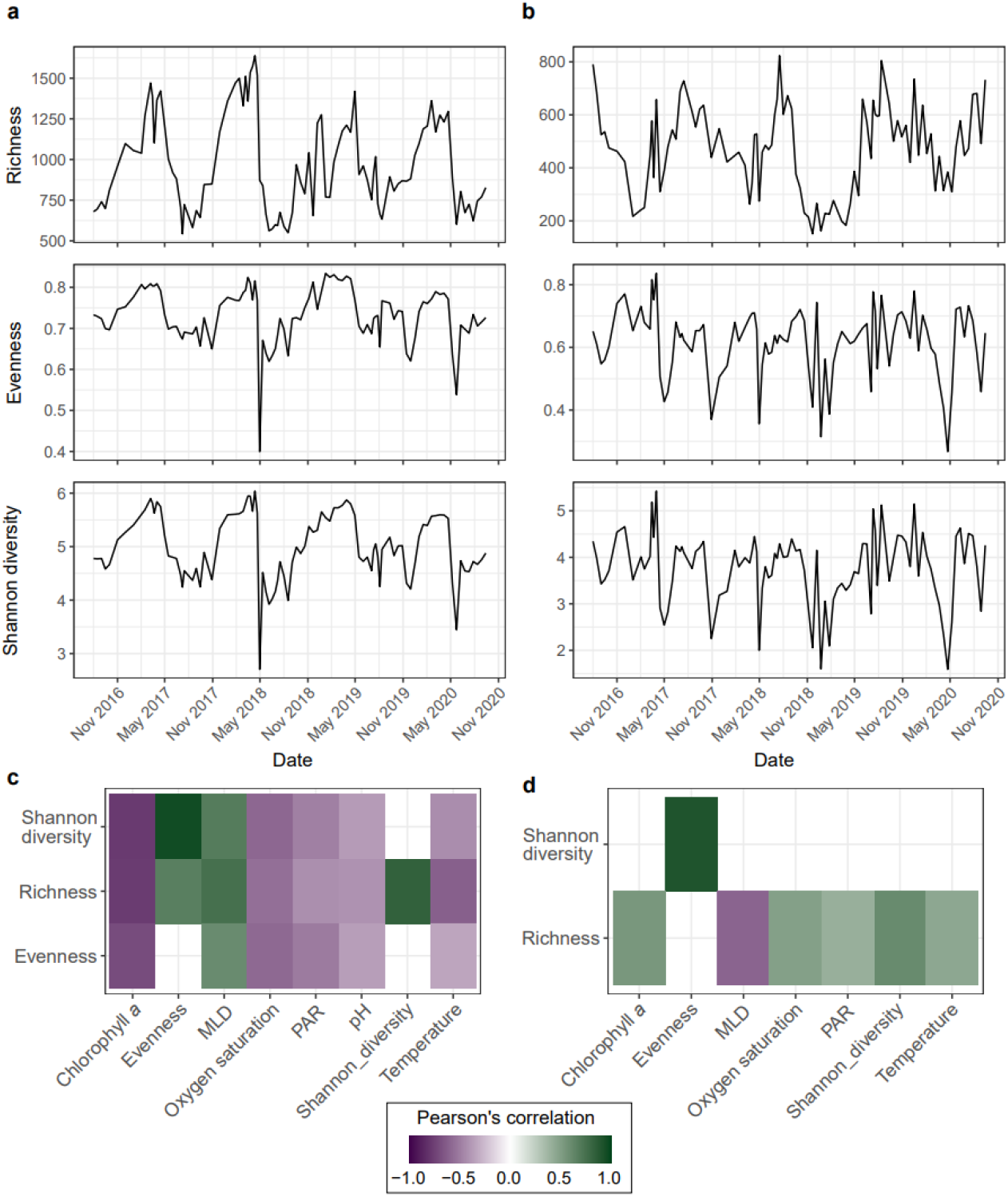
The diversity of prokaryotic and microeukaryotic communities is structured differently over time. All values of diversity shown represent the mean value after performing 100 iterations of rarefying and metric calculation. Richness, Evenness and Shannon diversity of **a)** prokaryotic and **b)** microeukaryotic communities. Statistically significant (p < 0.05) correlations between **(c)** prokaryotic and **(d)** microeukaryotic alpha-diversity measures and environmental parameters after multiple testing correction.

In contrast to prokaryotes, microeukaryotic communities exhibited a bimodal fluctuation in alpha diversity in each annual cycle. The bimodal pattern was reflected in a peak in *H’* in both polar-night (February-March) and polar-day (July-August). However, during polar night, the increased *H’* was underpinned by a reduced R and increased E, while in polar day, it was driven by an increased R. These temporal fluctuations in microeukaryotic diversity are in contrast to observations from the temperate San Pedro time-series (SPOT), where *H’* was shown to be invariable over time across the top 500 m of the water column^13^. The difference may indicate more seasonal structuring of microeukaryotic communities in high-latitude ocean ecosystems as a consequence of the pronounced environmental variability, supported by the negative correlation of MLD and Richness (Pearson’s *R* = −0.48, 95% CI [−0.62,−0.31], *p* = 7.2 × 10^−7^). Furthermore, the distinct trends observed between prokaryotes and microeukaryotes provides evidence that the diversity of these communities is shaped by different forces.

We next assessed how the composition of communities was structured over time. We observed a coherent structuring of prokaryotic and microeukaryotic communities based on the month of sampling. That is, communities sampled from the same month across years were often more similar than from other months in the same year (Figure 3a and 3b), reflecting an annual ecosystem clock. For prokaryotic communities, this pattern aligns with previous observations of month-based clustering at the Banyuls Bay Microbial Observatory in the Mediterranean^31^ as well as lowest pairwise beta-diversity at 12-month intervals reported from the temperate English Channel L4^32^ and SPOT^13^ time-series. However, in contrast, microeukaryotic communities showed weaker temporal structuring over multi-annual scales in the SPOT time-series^13^. By comparing the within- and between-month dissimilarities across years through the convex hull areas within the NMDS ordination (Figure 3 and Supplementary Table S6), we demonstrate a clear distinction in the temporal recurrence of prokaryotic and microeukaryotic composition. Prokaryotic communities were more cohesive across years in February-March and more variable during June-July, with convex hull areas of ∼0.025 and ∼0.13 respectively. In contrast, microeukaryotic communities were more cohesive during August and more variable during January-March, with convex hull areas of 0.06 and ∼0.43 respectively. However, maximal inter-annual differences in microeukaryotic communities were observed in April, in the phase before the spring bloom. This indicates that the assembly of the spring bloom is less predictable, and only later in the summer, the increased richness of microeukaryotic populations (high *R*) assembles into a cohesive structure each year. In previous years, a high inter-annual variability during polar day has been observed in microeukaryotic communities in this region^33,34^, so this pattern may change with time and reflect climate-induced or natural decadal variations^34^. The recurrent structuring of microeukaryotic communities during the productive season could be anticipated to stimulate predictable dynamics in prokaryotes, owing to specialised substrate niches and specific interactions^35,36^ as observed in temperate systems^7,8^. Indeed, it triggers a pronounced response of a minority of prokaryotic populations, evidenced through reduced *R* and *E*, but their emergent population structure shows high inter-annual variability, which may be a result of selection on a functional level and stochastic processes. In contrast, polar night conditions manifest species-rich but compositionally cohesive prokaryotic communities, suggesting that vertical mixing drives replenishment back to a “standing stock”.

**Figure 3.**
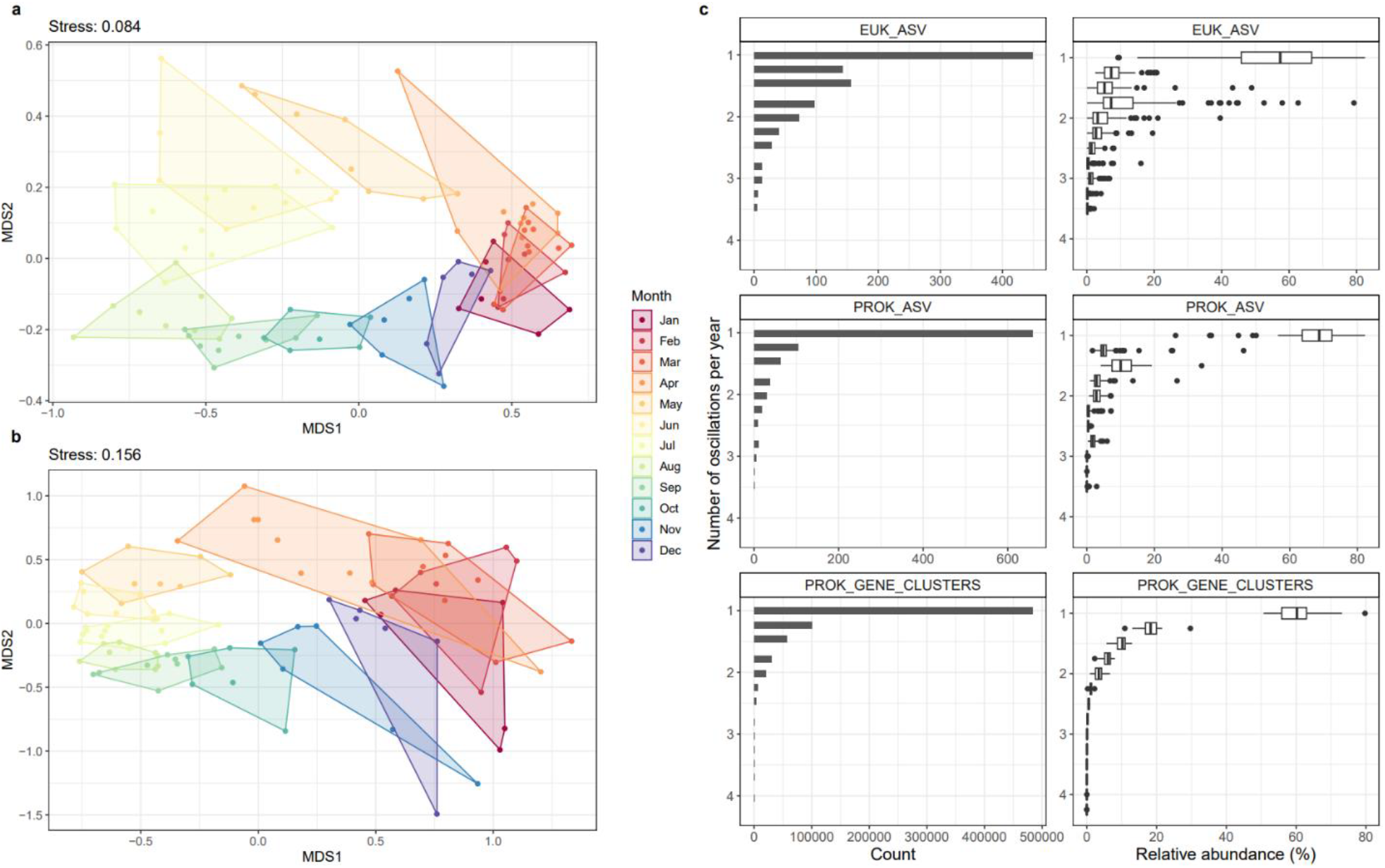
Structuring of community composition and the oscillations of ASVs and genes across years. Non-metric multi-dimensional scaling of Bray-Curtis dissimilarities calculated after rarefying and Hellinger transforming ASV count data for **(a)** prokaryotic and **(b)** microeukaryotic communities over four years, where colours indicate months. **c)** The count (left) and relative abundance (right) of prokaryotic ASVs, microeukaryotic ASVs, and prokaryotic gene clusters for each oscillation signal. The relative abundances of ASVs and genes was subject to Fourier transformation and oscillation signals determined based on amplitude of peak/trough dynamics within each annual cycle. An oscillation of 1 indicates a single peak/trough, reflecting a unimodal annual fluctuation in abundance.

#### Temporal dynamics of ASVs and community gene content

To gain a deeper understanding of ecosystem dynamics in the WSC, we complemented the amplicon dataset with long-read metagenomes and systematically investigated the temporal fluctuations of ASVs and community gene content. From 47 PacBio HiFi read metagenomes that spanned the four years, we obtained 48.5 million open-reading frames that could be classified as bacterial or archaeal, by comparing to species-representative genomes in the GTDB. The predicted prokaryotic gene sequences were subsequently grouped into 704,158 non-singleton clusters based on a 95% average nucleotide identity cut-off. This, combined with the ASV data, forms a rich multi-year ecosystem catalogue that can be used to elucidate community assembly processes and temporal dynamics on a taxonomic and functional level.

To unravel the temporal dynamics of ASVs and gene clusters, we employed an approach based on Fourier transformations and the determination of oscillation signals. Fourier transformations convert abundance data into frequencies, resulting in wave-like signals that can be evaluated in terms of peak/trough dynamics, hereon termed oscillation signals. We determined that 18% of prokaryotic ASVs, 15% of microeukaryotic ASVs and 69% of gene clusters exhibited a single oscillation each year, reflecting a unimodal fluctuation in abundance with a single peak and trough (Figure 3c). Although only capturing a fraction of the diversity, these annually oscillating ASVs and gene clusters comprised the majority portion of communities across all time points, with an average relative abundance of 67% for prokaryotic ASVs, 55% for microeukaryotic ASVs, and 60% for gene clusters. These findings expand on previous reports of seasonally recurrent patterns in temperate prokaryotic communities at more coarse-grained resolutions, such as OTUs^10^ and higher taxonomic levels^13,22^, and are in line with those on microdiversity temporal dynamics from a coastal temperate region^8^ – suggesting comparable ecological forcing in high-latitude oceans.

The dominance of annually oscillating ASVs raises the question, “To what extent are these patterns deterministic?”. More than 15 years ago, Fuhrman and colleagues proposed that “annually recurrent microbial communities can be predicted from ocean conditions”^12^. However, only more recently, with technological advancements and increasing quantity and resolution of data, is predictive ecology becoming more feasible. To contribute to this, we assessed the timing and order of ASV oscillations across each annual cycle. We found that 20% of prokaryotic ASVs consistently oscillated in the same order each year, while 51% reached their peak within the same 30-day window. Therefore, while recurrent oscillations are largely bound within temporal windows, the composition of co-occurring populations can vary across years, likely reflecting the influence of trophic interactions and the ephemeral nature of population dynamics^14,37^. A similar pattern was recently described from prokaryotic communities in the NW Mediterranean, where seasonally recurrent taxa exhibited inter-annual changes in the composition of their neighbours within co-occurrence networks^31^. Despite these variations, the periodic timing of recurrent population dynamics within narrow temporal windows provides strong support towards deterministic patterns in ocean ecosystems.

#### Seasonal recurrence is underpinned by transitions across distinct ecological states

We investigated how the annual recurrence of ASVs and genes translates to ecological and functional shifts within prokaryotic communities. For this, we first grouped the prokaryotic genes into functions. Of the 482,923 annually oscillating gene clusters, 85% were assigned to an orthologous group in the EGGNOG database and were subsequently grouped based on the functional annotation of the matching ortholog. This resulted in 11,320 unique gene functions that captured between 41 – 72% of community gene content across all time points.

To investigate temporal structuring on a taxonomic and functional level, we used the oscillations of prokaryotic ASVs and gene functions to build a correlation-based network. Correlation co-occurrence networks have proven powerful for disentangling community dynamics and organismal interactions over spatial and temporal scales^38^. Here, we compared the oscillation signals of ASVs and gene functions through Pearson’s correlation and retained only those with a strong, positive coefficient (*R* > 0.7, FDR-based adjusted *p* < 0.05). We subsequently built a network using these correlation coefficients as the edges and the respective ASVs and gene functions as the nodes. Using the Louvain algorithm, the network was partitioned into five modules comprised of co-oscillating ASVs and gene functions (Figure 4a) that represent distinct temporal periods in the WSC. Each annual cycle was thus characterised by a succession across these five temporal modules (Figure 4b and Supplementary Table S7).

**Figure 4.**
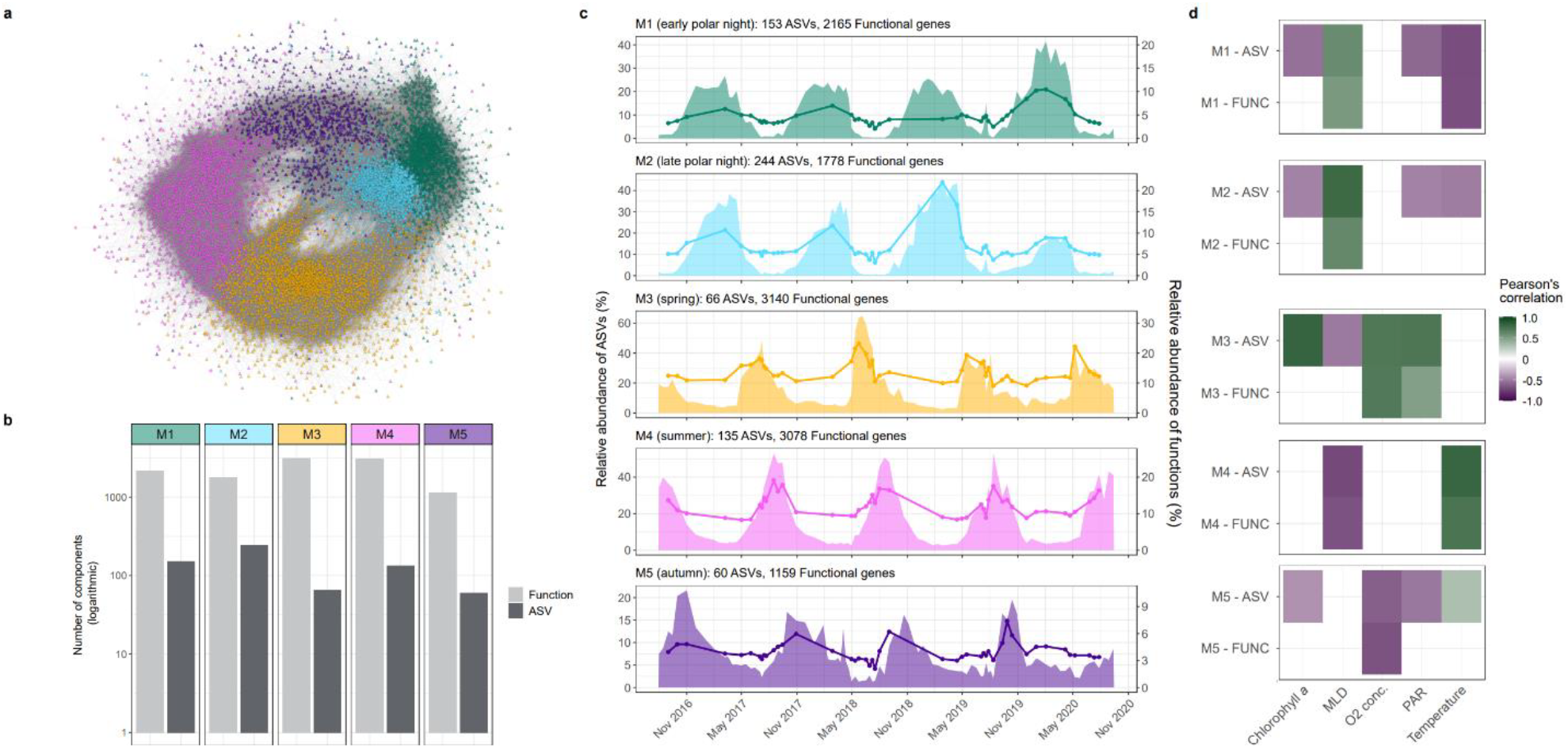
Ecosystem modules, their temporal dynamics and association with environmental conditions. **a)** Co-occurrence network constructed from significant positive Pearson correlations (>0.7) between prokaryotic ASVs and functional cluster oscillation signals. Circle nodes = ASVs, Triangle nodes = functional clusters. **b)** The temporal relative abundance dynamics of module ASVs (area) and functional clusters (line). **c)** Significant Pearson’s correlations between the combined relative abundance of module ASVs and module functional clusters against measured environmental factors (only those with a p <0.05 after multiple testing correction are shown).

To uncover the ecological shifts associated with the module succession, we compared the composition of modules, the environmental conditions they prevail under, and their association with microeukaryotes. The modules differed markedly in the number of their taxonomic and functional components, from a low number of ASVs (n = 66) and high number of functions (n = 3140) in the spring module M3 to a low number of both functions (n = 1159) and ASVs (n = 60) in the autumn module M5. Beyond quantitative differences, the modules comprised unique taxonomic (Figure 5a) and metabolic signatures (Figure 5b). Module M1, which prevailed from early to mid-polar night, was taxonomically distinguished by *Cand*. Nitrosopumilus, Arctic97B-4 (Verrucomicrobiota) and BD2-11 (Gemmatimonadota) and functionally distinguished by ammonia oxidation (*amoABC*), nitrite reduction (*nirK* and *norBC*) and carbon fixation (hydroxypropionate/hydroxybutyrate cycle^39^). The numerous *Cand*. Nitrosopumilus ASVs detected and their high network connectivity to gene functions within module M1 (Figure 5a) complements previous findings from southward flowing Arctic waters^40^ and reports of increased ammonia oxidation rates during polar night in Antarctic coastal waters^41^. Taken together, these observations indicate that ammonia-oxidising Archaea are likely keystone members of prokaryotic communities during winter in high-latitude ocean ecosystems, irrespective of water mass origin. Microeukaryotic ASVs that co-oscillated with this early polar night module included members of Radiolarians, Nassophorea and MOCH-1 (Figure 6). A shift in dominance from module M1 to M2 marked a transition in the prevailing ecological state during the mid- to late-polar night period. This transition saw the emergence of *Thalassobius* and *Arenicellaceae* members dominating the prokaryotic communities along with an increase in C1 and nitrogen metabolic machinery. The onset of solar radiation and rapid transition from module M2 to M3 signified the end of polar night. The positive correlation of module M3 with PAR and chlorophyll (Figure 4c and Supplementary Table S8) as well as its association with Bacillariophyta, Prymnesiophyceae and Dinophyceae (Figure 6) indicates its representation of the spring phytoplankton bloom period. As could be expected, this module was dominated by members of heterotrophic bacteria known as primary responders to phytoplankton blooms and their carbohydrate exudates in temperate regions^7,35^, including *Polaribacter, Aurantivirga, Formosa* and SAR92. Functionally, module M3 was enriched in carbohydrate-active enzymes, amino and nucleotide sugar metabolism and organosulfur compound utilisation, including DMSP demethylation, DMSO to DMS (*dmsBC*) and methanesulfonate to sulfate (*msmAB, sorAB*/SUOX sulfite oxidase). The summer module M4 that preceded the spring bloom was also enriched in sulfur metabolism but included both inorganic and organic sulfur utilisation, such as sulfur oxidation (*soxABCXYZ*), sulfite reduction (*soeABC*), DMSP demethylation, methanethiol oxidation (*MeSH*), and taurine utilisation (*tauABC*). The prokaryotic community in module M4 was dominated by a single *Amylibacter* ASV. *Amylibacter* are key contributors to organosulfur compound metabolism in temperate coastal waters during summer^42,43^. However, we could also attribute *Amylibacter* to the machinery to oxidise sulfur (SOX system) and reduce sulfite (sulfite reducatese), suggesting their specialisation on more diverse sulfur sources^44^. Module M4 was also signified by motility machinery, quorum sensing and photosynthesis. The photosynthesis machinery was associated with *Synechococcus*, which represented up to 1.7% of the prokaryotic community during this period. *Synechococcus* is a significant contributor to primary production in tropical and temperate oceanic regions but is largely absent from polar waters. Our observations of increased *Synechococcus* abundances in late summer mirrors recent findings from the WSC^45^ and suggests that progressing Atlantification is driving their northward expansion into the Arctic.

**Figure 5.**
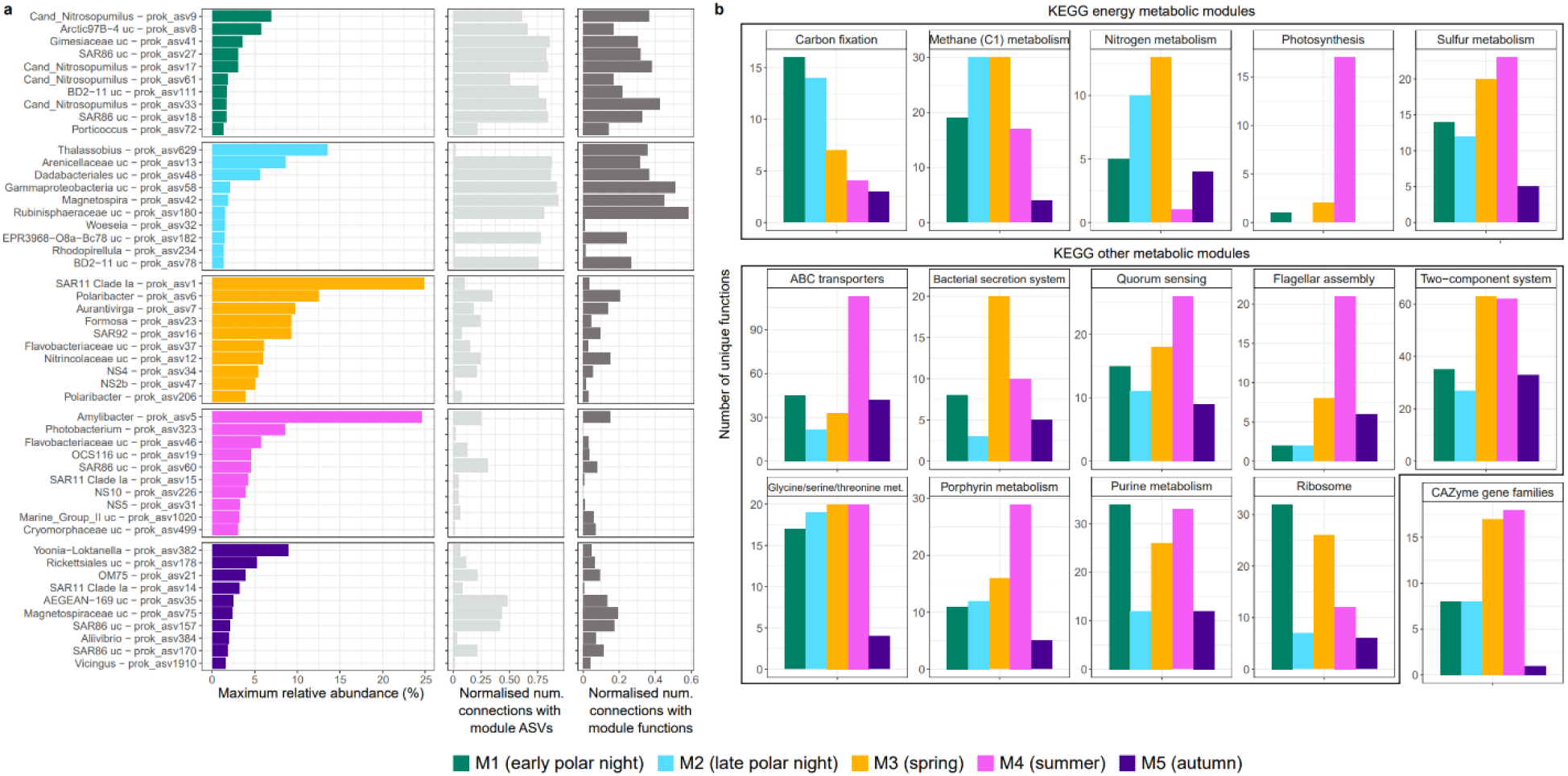
Modules are phylogenetically and functionally distinct. **a)** Ten most abundant ASVs identified in each module, along with the number of connections they share with other ASVs and functions from the same network module. The number of network connections was normalised by the total number of nodes in each module. **b)** Composition of KEGG metabolic pathways that exhibited the largest variance in the number of assigned functions between modules along with the number of carbohydrate-active enzyme gene families.

**Figure 6.**
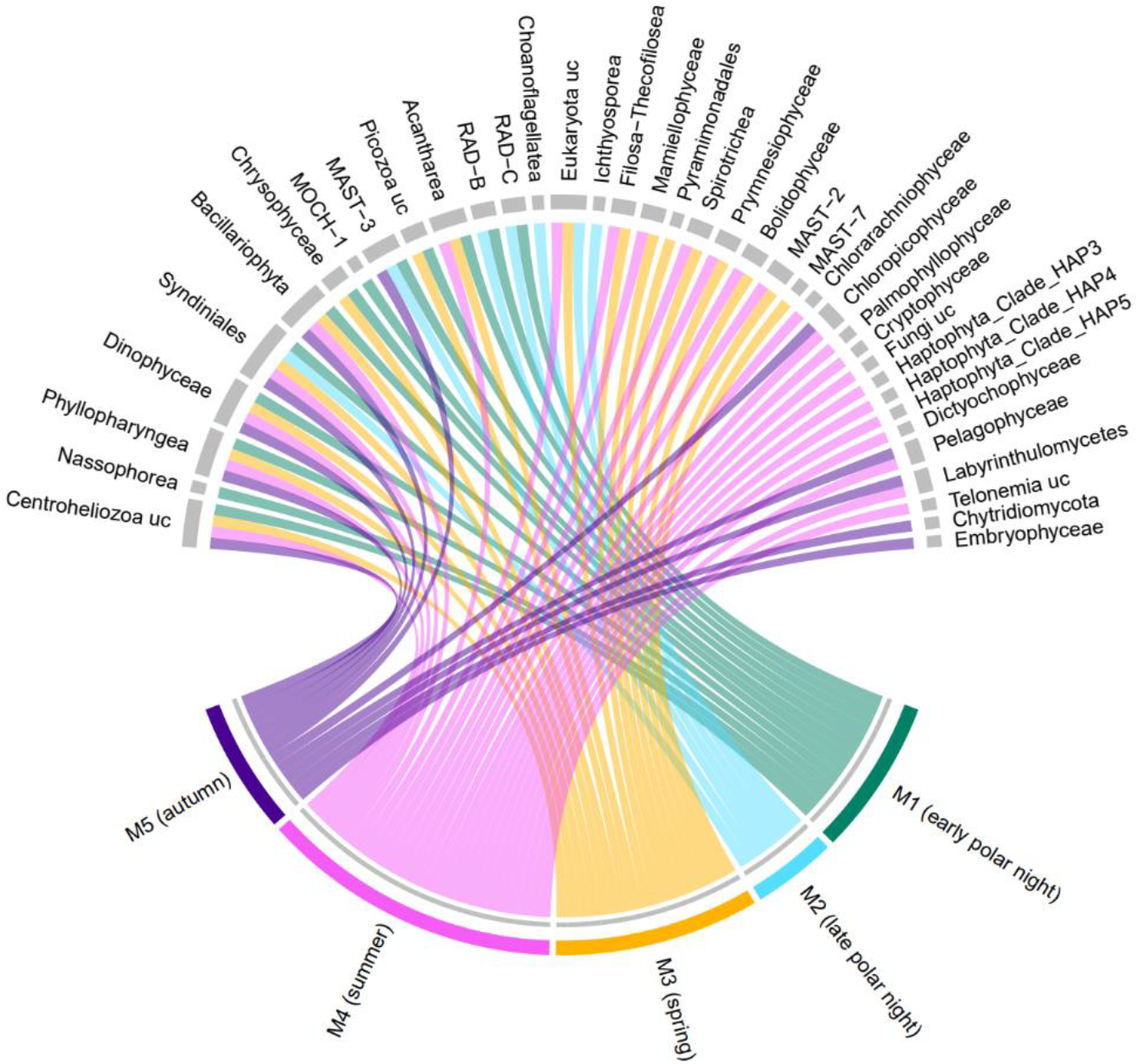
Correlations between prokaryotic and microeukaryotic ASV oscillations. Pairwise Pearson’s correlations were computed between module associated prokaryotic ASVs and microeukaryotic ASVs. Correlations with a coefficient > 0.7 and p <0.05, after multiple testing correction, were used to construct the chord diagram. Microeukaryotic ASVs are grouped at higher taxonomic ranks.

The taxonomic and metabolic signatures of each module demonstrate that the prokaryotic microbiome is structured by a succession across five distinct ecological states within each annual cycle. Previous taxonomic-centred analyses of prokaryotic communities from temperate and tropical ecosystems have also reported recurrent dynamics structured by seasonality. However, the proportion of ASVs exhibiting seasonal recurrence and the number of temporal modules appears to be higher in the WSC. For instance, in the North West Mediterranean, only 4% of prokaryotic ASVs, constituting a relative abundance of 47%, were shown to exhibit seasonal recurrence and could be grouped into three distinct seasonal clusters^46^. In a coastal temperate region that lacks pronounced phytoplankton blooms but experiences large environmental variability, recurrent dynamics of prokaryotes are primarily partitioned into summer and winter groups^47^. The higher prevalence of recurrent dynamics and their organisation into narrower temporal modules in the WSC may indicate stronger selective pressure arising from the more pronounced seasonal environ mental variability.

#### Selection pressure varies across ecological states

The recurrent dynamics of ASVs, genes and functions and their assembly into cohesive, ecological modules suggests that the WSC microbiome is predominantly shaped by deterministic processes. The primary mechanisms that contribute to driving deterministic structuring of microbiomes are environmental selection and organismal interactions. Environmental selection is considered the primary force shaping the distribution of populations and the assembly of microbial communities in ocean ecosystems^48^. However, we lack a mechanistic understanding of how environmental selection operates. In particular, it is unclear whether the environment primarily selects for a function or for a specific organism with a function. Recent evidence has shown that the taxonomic composition of a microbiome can change while the gene content remains conserved^49,50^, reflecting the prevalence of metabolic redundancy across microbial taxa. In addition, studies on surface ocean microbial communities have demonstrated incongruencies in structuring on a taxonomic and functional level, with strong selection pressure on functional groups but weak selection pressure on the taxonomic composition within functional groups^18^. These observations provide evidence for the decoupling of taxonomy and function within microbiomes and suggests that selection may act on a functional level.

Here, we showcase that selection acts heterogeneously across seasonal periods, with differential pressures on a taxonomic and functional level. To demonstrate this, we focus on the late polar night module M2 and spring module M3, as they represent two contrasting scenarios. The spring module M3 comprises a nearly two-fold larger diversity of gene functions than the polar night module M2, but fourfold less ASVs. Hence, spring is underpinned by annually recurrent dynamics predominantly on a functional level, compared to both taxonomic and functional recurrence during late polar night. Hence, selection pressure may be stronger on the function than organismal level in spring. However, despite this, both modules contained more functional redundancy than the other seasonal modules, with 10% more multi-gene cluster compared to single-gene cluster functions. Amongst these redundant functions, we identified a strong positive linear relationship between function abundance and the diversity of the contained gene clusters (Figure 7b) in ∼55% of cases. As such, the oscillations of these redundant functions are, in part, driven by the concurrent oscillations of metabolically overlapping prokaryotes as opposed to selection for specific populations. To explore this further, we compared the structuring of gene cluster diversity within redundant functions over time. In module M2, 60% of the redundant functions were dominated by the same gene cluster during each annual oscillation, compared to 40% in module M3 (Figure 7c). The increased genetic variability within functions of module M3 supports the notion of stronger functional selection and weaker organismal selection during spring.

**Figure 7.**
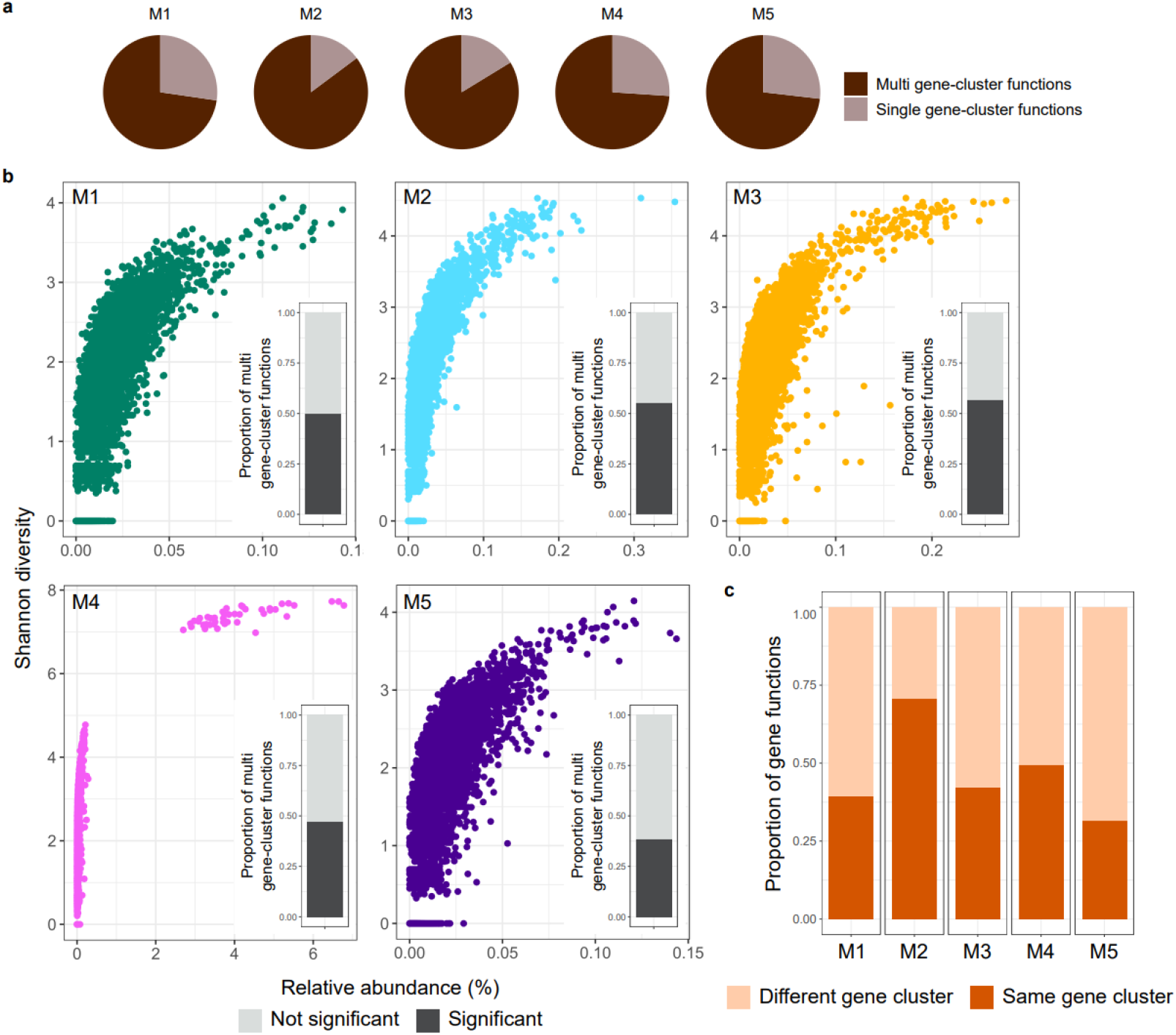
Diversity, abundance and structuring of gene clusters within functions. **a)** Proportion of functions in each module comprised of multiple or single gene clusters. For those functions that contain >1 gene cluster, we calculated the **b)** Shannon diversity of gene clusters within functions against the relative abundance of the function across all timepoints. Each point represents an individual function. The inserted barplots illustrate the proportion of multi gene clusters functions that exhibit a significant positive linear relationship between Shannon diversity and abundance. **c)** Proportion of multi gene clusters functions that are dominated by the same or different gene cluster during each annual oscillation.

Integrating the observations on ASV dynamics further supports seasonal variations in environmental selection pressure. The late polar night features high species richness and evenness and a low variability in community composition across years. As such, the late polar night is characterised by a well-mixed community that is taxonomically and functionally conserved across years. Interestingly, we observed three ASVs during this period that reached a large fraction of the microbial community (Figure 5), which included members of known sub-surface taxa, such as *Dadabacteriales*. Therefore, the deeper vertical mixing during polar night drives a recurrent homogenisation of epipelagic communities, with potentially a weaker environmental pressure that selects for only a few populations upwelled from subsurface waters. Conversely, spring exhibits low species richness and evenness and high inter-annual variability in community composition (Figure 2 and 3). Combined with the observations on gene functions, spring can thus be characterised by the emergence of a prokaryotic community dominated by few populations that can compositionally vary but remain functionally conserved across years. Therefore, selection pressure appears stronger on a functional rather than organismal level in spring.

To further advance our understanding on environmental selection and the mechanisms that shape the structuring of ocean microbiomes, it is paramount to integrate both taxonomic and functional profiling along with more extensive environmental information. Of particular importance is information on the availability of organic carbon and energy substrates, which play a fundamental role in shaping the dynamics of microbial communities. For instance, phytoplankton blooms in coastal temperate regions that are of a larger magnitude than in the WSC have been shown to drive a predictable composition of microbes^7,8^, in contrast to our observations here. The disparity between the regions may be related to the quantity of organic matter released by such phenomena, i.e. higher organic matter concentrations may drive more deterministic taxonomic responses. However, whether the availability of organic matter, or other here unmeasured environmental factors, play a role in shaping the deterministic and recurrent dynamics within microbiomes requires further investigation.

## CONCLUSION

Our study provides fundamental insights into the seasonal and interannual structuring of prokaryotic and microeukaryotic communities in an Arctic pelagic ocean ecosystem. We demonstrate the prevalence of annually recurrent dynamics of populations and community gene content, which are organised into five distinct seasonal modules. Each of the modules represents distinct ecological states that prevail within the prokaryotic microbiome each year and are connected to specific microeukaryotic populations and environmental conditions. We further provide evidence that environmental selection is heterogeneous across these ecological states, with differential pressures on an organismal and functional level. Our findings provide new insights into understudied yet rapidly changing ocean regions and advances our understanding of how microbial communities are structured across pronounced environmental gradients.

## MATERIALS AND METHODS

### Sample collection and processing

Moorings carrying autonomous water samplers (Remote Access Samplers; RAS) were deployed between 2016 – 2020 at a single location in the eastern Fram Strait (mooring F4: 79.0118 N 6.9648 E). Moorings were deployed for 12-month intervals, with collection and redeployment occurring in summer, typically August. Owing to ocean currents, the vertical positioning of the RAS fluctuated between 20 – 110 m over the four-year period. At weekly to fortnightly intervals, 2 × 500 ml of seawater was collected in sterile plastic bags and fixed with mercuric chloride (0.01% final concentration). Following mooring recovery, fixed seawater samples from each timepoint were filtered onto 0.22 μm Sterivex cartridges and directly frozen at −20 °C until DNA extraction.

### Mooring and satellite data

Attached to the RAS were Seabird SBE37-ODO CTD sensors that measured temperature, depth, salinity, and oxygen concentration. Sensor measurements were averaged over 4 h around each seawater sampling event. Physical sensors were manufacturer-calibrated and processed in accordance with https://epic.awi.de/id/eprint/43137. Employing multiple CTD sensors along the mooring depths enabled the determination of the minimum mixed layer depth (MLD) at each sampling time point. For instance, if two CTDs showed the same temperature and salinity measurements, the MLD was at least the depth of the deeper CTD. Chlorophyll concentrations were measured via Wetlab Ecotriplet sensors. Surface water Photosynthetically Active Radiation (PAR) data, with a 4 km grid resolution, was obtained from AQUA-MODIS (Level-3 mapped; SeaWiFS, NASA) and extracted in QGIS v3.14.16 (http://www.qgis.org).

### SSU rRNA gene amplicon and metagenome sequencing

Filtered seawater samples from 97 time points were subjected to DNA extraction using the DNeasy PowerWater Kit (QIAGEN, Hilden, Germany). 16S and 18S rRNA gene fragments were PCR-amplified using the primers 515F–926R^51^ and 528iF–964iR^52^, respectively. Sequencing libraries were constructed from rRNA gene products according to the “16S Metagenomic Sequencing Library Preparation” protocol (Illumina, San Diego, CA) and sequenced on an Illumina MiSeq platform in 2 × 300 bp, paired-end mode. Amplicon sequencing took place at the Alfred Wegener Institute. The extracted DNA from 47 timepoints was additionally used to generate PacBio HiFi metagenomes. Sequencing libraries were prepared following the protocol “Procedure & Checklist – Preparing HiFi SMRTbell Libraries from Ultra-Low DNA Input” (PacBio, Menlo Park, CA) followed by inspection with a FEMTOpulse. The libraries were multiplexed and sequenced on 8M SMRT cells (7 - 8 samples per cell) on a PacBio Sequel II platform for 30 h with sequencing chemistry 2.0 and binding kit 2.0. Metagenomes were sequenced at the Max Planck Genome Centre, Cologne, Germany.

### PacBio HiFi metagenome analysis

A total of 48 Gbp of PacBio HiFi reads were generated, with an average of 1 Gbp per sample. Gene sequences were predicted on HiFi reads using Fraggenescan (v1.31; parameters:-complete=1 - train=sanger_5)^53^. The genes were subsequently clustered using cd-hit v4.8.1^54^ at a 95% identity threshold – these comprise the ‘gene clusters’. To facilitate comparisons between metagenomes, gene cluster counts were normalised by the estimated number of prokaryotic genomes in each sample, determined from the average sequencing depth of 16 single-copy ribosomal proteins, as described previously^55^. The longest sequence from each cluster was used as the representative for functional assignment against the EGGNOG v5.0 database^56^ using the eggnog-mapper tool v2^57^. To build the ‘functional clusters’ used in our analysis, we grouped gene clusters based on functional annotations of matching seed orthologs. As the seed orthologs of EGGNOG have been functionally annotated using numerous resources, we grouped gene clusters firstly based on annotations against the carbohydrate-active enzyme database, followed by KEGG and then PFAM.

### Taxonomic diversity analyses

The 16S and 18S amplicons were processed into Amplicon Sequence Variants (ASV) using DADA2^58^ in RStudio v4.1.3^59^. ASVs were taxonomically classified using the SILVA SSU v138 (16S) and PR2 v4.12 (18S) databases using the *assignTaxonomy* function of DADA2. The ASV dataset was filtered to remove those with <3 counts in <3 samples. For alpha diversity, the ASV count table was subject to 100 iterations of rarefying followed by the calculation of Richness, Shannon diversity (*diversity* function in *vegan*) and Evenness. The mean and standard deviation was calculated for each metric, with the mean being used for visualisations and statistical comparisons to environmental metadata. For beta diversity analysis, the ASV count table was rarefied a single time, followed by Hellinger transformation before calculation of Bray-Curtis dissimilarities (using *vegan*). Dissimilarities were ordinated using Non-Metric Multi-Dimensional Scaling.

### Time-series and network analysis

The temporal analysis of ASVs and gene clusters as well as the construction of co-occurrence networks was performed using the same workflow as that presented recently by authors of this paper^60^. In brief, for each ASV and gene cluster, we calculated a time-series signal using the signal using the *segmenTier*/*segmenTools* (https://github.com/raim/segmenTools) packages https://www.nature.com/articles/s41598-017-12401-8) using relative (ASV) and normalized (gene cluster) abundance data. The recovered signals indicate the frequency, amplitude and phase of abundances in each annual cycle, which we termed oscillation signals. Oscillation signals were extracted for each ASV and gene cluster using the *fprocessTimeseries* function from the *segmenTier* package. The ASVs and genes with an oscillation signal of one, i.e. a single peak and trough in each annual cycle, were deemed as “annually oscillating” and retained for further analysis. Phase-Rectified Signal Averaging (PRSA) was used to visualize periodic patterns of ASVs, using phase-rectified data to remove phase variability. Then, the phase-rectified data was averaged for the final PRSA plot. The oscillation signals of ASVs and gene clusters were compared through pairwise Pearson’s correlation, with multiple testing corrections using the FDR method. Those with a statistically significant (*p* < 0.05) positive correlation coefficient of >0.7 were used to build a co-occurrence network, with edges as correlation coefficients. By using oscillation signals, focusing on ASVs and gene clusters with a defined oscillation, and using only positive correlations, we minimise noise in the dataset and prevent potential network topology distortion of negative correlations. Co-occurrence correlation networks were constructed using the *igraph* package^61^ in R and visualised in Cytoscape v3.7.2^62^ using the Edge-weighted Spring-Embedded Layout. ASVs and gene clusters in the network were clustered using the Louvain algorithm^63^. All steps outlined above were performed in R v4.1.3.

## Supporting information

Supplementary tables all

## DATA AVAILABILITY

Mooring data are available under https://doi.pangaea.de/10.1594/PANGAEA.904565 (2016-2017), https://doi.pangaea.de/10.1594/PANGAEA.904534 (2017-2018), https://doi.pangaea.de/10.1594/PANGAEA.941126 (2018-2019), and https://doi.pangaea.de/10.1594/PANGAEA.946508 (2019-2020). ENA accession numbers for 16S rRNA amplicons are: PRJEB43890 (2016-2017), PRJEB43889 (2017-2018), PRJEB67813 (2018-2019), PRJEB66202 (2019-2020). ENA accession numbers for 18S rRNA amplicons are: PRJEB43504 (2016-2017), PRJEB43885 (2017-2018), PRJEB66212 (2018-2019), PRJEB66220 (2019-2020). Raw metagenomic reads are available under PRJEB67368. Code for reproducing workflow and figures is available at https://github.com/tpriest0/Fram_Strait_WSC_time_series_2016-2020 while the accompanying derived data files are available at https://doi.org/10.17617/3.CA8MQY.

## ACKNOWLEDGEMENTS

We thank Jana Bäger, Theresa Hargesheimer, Rafael Stiens and Lili Hufnagel for RAS operations; Daniel Scholz for RAS and sensor operations; Normen Lochthofen, Janine Ludszuweit, Lennard Frommhold and Jonas Hagemann for mooring operations; Jakob Barz, Swantje Ziemann and Anja Batzke for DNA extraction, library preparation and sequencing; and Bruno Huettel and the technicians at the Max Planck Genome Centre in Cologne for sequencing. The captain, crew and scientists of RV Polarstern cruises PS99.2, PS107, PS114, PS121 and PS126 are gratefully acknowledged. This project has received funding from Polarstern grants AWI_PS99_00, AWI_PS107_05, AWI_PS114_01, AWI_PS121_07, AWI_PS126_05, and AWI_PS126_07. Further support came from the European Research Council (ERC) under the European Union’s Seventh Framework Program (FP7/2007-2013) research project ABYSS (Grant Agreement no. 294757) to AB, from the Helmholtz Association, specifically for the FRAM infrastructure, and from the Max Planck Society.

## AUTHOR CONTRIBUTIONS

TP designed and executed the ASV and metagenomic analysis pipelines and wrote the manuscript, with input from MW, EO and BD. EO and OP performed time-series and network analyses. MW processed amplicon raw data into ASVs, co-designed the sampling and mooring strategy, and coordinated data analysis. WJvA contributed quality-controlled oceanographic data and coordinated the mooring operations. KM coordinated the processing of samples and sequencing and provided 18S rRNA gene sequence data. STV provided quality-controlled chlorophyll sensor data. CB, KM and AB co-designed the sampling and mooring strategy, and contributed to the interpretation. BF and RA contributed to study design and interpretation. All authors contributed to the final manuscript.

